# A comparative study of algorithms detecting differential rhythmicity in transcriptomic data

**DOI:** 10.1101/2023.10.12.562079

**Authors:** Lin Miao, Douglas E. Weidemann, Katherine Ngo, Benjamin A. Unruh, Shihoko Kojima

## Abstract

Rhythmic transcripts play pivotal roles in driving the daily oscillations of various biological processes. Genetic or environmental disruptions can lead to alterations in the rhythmicity of transcripts, ultimately impacting downstream circadian outputs, including metabolic processes and even behavior. To statistically compare the differences in transcript rhythms between two or more conditions, several algorithms have been developed to analyze circadian transcriptomic data, each with distinct features. In this study, we compared the performance of seven algorithms that were specifically designed to detect differential rhythmicity. We found that even when applying the same statistical threshold, these algorithms yielded varying numbers of differentially rhythmic transcripts. Nevertheless, the set of transcripts commonly identified as differentially rhythmic exhibited substantial overlap among algorithms. Furthermore, the phase and amplitude differences calculated by these algorithms displayed significant correlations. In summary, our study highlights a high degree of similarity in the results produced by these algorithms. Furthermore, when selecting an algorithm for analysis, it is crucial to ensure the compatibility of input data with the specific requirements of the chosen algorithm and to assess whether the algorithm’s output fits the needs of the user.

## Introduction

Most living organisms exhibit circadian rhythms, which are internal processes that drive biochemical and behavioral changes approximately every 24 hours (1). The rhythmicity of these processes is driven ultimately by rhythmically expressed transcripts, which are regulated by the molecular clock machinery (2-8). Genetic or environmental disruptions can lead to alterations in the rhythmicity of these transcripts, ultimately impacting downstream circadian outputs, including metabolic processes and behavior (9-15).

To detect differences in rhythms between groups or conditions in transcriptomic datasets, different approaches have been used. One option is Venn Diagram Analysis (VDA). In this approach, transcripts are independently categorized as either rhythmic or non-rhythmic under each condition using a rhythmicity detection algorithm with an arbitrary statistical threshold (16-20). Subsequently, transcripts that are categorized differently between conditions are recognized as differentially rhythmic. This approach offers a straightforward means of distinguishing transcripts that are rhythmic under one condition but not under the other. However, it presents several challenges: 1) the analysis is indirect; no statistical test is performed to reject the null hypothesis that there are no differences in rhythmicity between conditions, 2) this approach can only identify transcripts that are rhythmic in one condition but not in the other and does not detect quantitative changes in their rhythms (e.g., changes in amplitude or phase), and 3) this approach generally overestimates the number of differentially rhythmic transcripts because of the uncontrolled false discoveries from two individual rhythmicity tests (21-23).

More recently, several algorithms have been developed to statistically compare rhythmicity between two or more conditions. Even though these algorithms share the same goal, their approaches in detecting rhythmic transcripts and differentially rhythmic transcripts are different. In addition, the requirements for input datasets and the output information are also often different. Here, we focus on seven of these algorithms published or posted on bioRxiv in or before 2022: DODR (22), LimoRhyde/LimoRhyde2 (24, 25), CircaCompare (26), compareRhythms (21), diffCircadian (23), dryR (27), and RepeatedCircadian (28), and perform a systematic comparison of their characteristics and performance. We highlight three features of these algorithms: 1) the requirements for input data, 2) the number of rhythmic and differentially rhythmic transcripts detected, and 3) the differences in their outputs. We also analyzed the same RNA-seq dataset with each approach in order to directly compare their performance. We uncovered both notable differences and similarities across these algorithms and our study will help guide future users in choosing the algorithm that best suits both the user’s dataset and the desired output.

## Materials and Methods

All analyses were performed in R (version 4.3.1), and all graphs were created with the ggplot2 (version 3.4.2), upsetR (version 1.4.0), and corrplot (version 0.92) R packages. Pearson’s correlation analysis was performed using the cor() function from Hmisc package (version 5.1-0). The circular version of Pearson’s correlation analysis was performed using the cor.circular() function from circular package (version 0.4-95).

### Input dataset

We used the RNA-seq dataset NIH SRA (Submission ID: 13164610) (9) for all analyses. Reads were mapped to the mouse mm10 genome (GENECODE:GRCm38.p6.genome.fa) and transcripts per million (TPM) and reads per kilobase per million mapped (RPKM) were quantified with HOMER (V4.11.1) (29) using the gencode.vM25.annotation.gtf with the option of condenseGenes. Transcripts with mean TPM < 0.25 (averaged across all time points and conditions) were removed from the analysis. Rhythmicity of RPKM was assessed by MetaCycle (v 1.2.0) (19), which was run using the following parameters: minper = 20, maxper = 28, and cycMethod = c(“ARS”, “JTK, “LS”). Transcripts with Benjamini-Hochberg adjusted q values (B.H. q) < 0.25 were considered rhythmic.

### Differential Rhythmicity Algorithms

The DODR R package (version 0.99.2) (22) was downloaded from https://cran.r-project.org/src/contrib/Archive/DODR/ and manually installed in R from the tar file, since this package is no longer available from CRAN. DODR was run with the RPKM data and used the following parameters: norm = TRUE, period = 24, and method = “robust”. The p-values provided by DODR were manually adjusted using the Benjamini-Hochberg procedure (30).

The LimoRhyde R package (version 1.0.1) (24) was run using the RPKM data with period = 24. The rhythmic transcripts under the control condition were identified using the limorhyde() function followed by limma R package (version 3.56.2) (31, 32) with the option of “trend = TRUE” for the function eBayes(), and the default options were used for other functions. The p-values were manually adjusted using the Benjamini-Hochberg procedure. The LimoRhyde2 R package (version 0.1.0) (25) was run using the RPKM data with the option of “sinusoid = TRUE” for the function of getModelFit(), and the default options were used for other functions.

The CircaCompare R package (version 0.1.1) (26) was run using the RPKM data with the default options, and the output p-values were manually adjusted using the Benjamini-Hochberg procedure. Transcripts with either significantly different amplitudes or phases were defined as differentially rhythmic. The circa_single() function was used to detect the rhythmicity of transcripts from the control condition.

The compareRhythms R package (version 0.99.3) (21) was run using the RPKM data with the options of method = “mod_sel”, “just_classify = FALSE”, and default parameters. Transcripts classified as having “gain”, “loss”, or “change” of rhythmicity were defined as differentially rhythmic.

The diffCircadian R package (version 0.0.0) (23) was run using the RPKM data with the options of period = 24, and method = “LR”. The p-values provided by diffCircadian were manually adjusted using the Benjamini-Hochberg procedure.

The dryR R package (version 1.0.0) (27) was run using the RPKM data with the drylm() function. Transcripts whose best fitting model was either 2 (rhythmic only in control), 3 (rhythmic only in *Bmal1* KD), or 5 (rhythmicity changed between the two conditions) were defined as differentially rhythmic. The dryseq_single() function was applied to detect the rhythmicity of transcripts from the control condition with untransformed count data because normalized data (e.g. RPKM, TPM) cannot be accepted.

## Results

### Characteristics of algorithms for detecting differential rhythmicity

We analyzed the seven algorithms that are the focus of this study and summarized their characteristics in Table 1. We first compared the input data requirements, as they differ in the types of data they can analyze (Table 1). Our focus centered on several critical characteristics including the algorithm’s ability to use untransformed count data (e.g. read counts), deal with uneven sampling intervals, handle missing samples or missing values for individual transcripts, and fit non-sinusoidal rhythm patterns. We also identified options to adjust the rhythm’s period and whether the algorithm required data with repeated measurements. We found that LimoRhyde2, compareRhythms, and dryR can accept untransformed count data, while all the other algorithms require normalized data (e.g., RPKM, CPM, or log-transformed expression values). All algorithms can accept datasets with uneven sampling intervals and missing one or more entire samples or time points. Except for compareRhythms and diffCircadian, the algorithms also accept missing values for individual transcripts in individual samples. Only LimoRhyde and LimoRhyde2 have the option to fit non-sinusoidal curves since these two algorithms have the option to fit both cosinor and periodic spline regression models. RepeatedCircadian is only compatible with datasets with repeated measurement experimental designs that have multiple measurements from the same sample over a time period (e.g., measurement of blood pressure at different time points within a circadian cycle of the same individual), and not measurements from different independent samples at each time point (e.g., datasets from liver samples collected from different mice at different times).

**Table 1.**
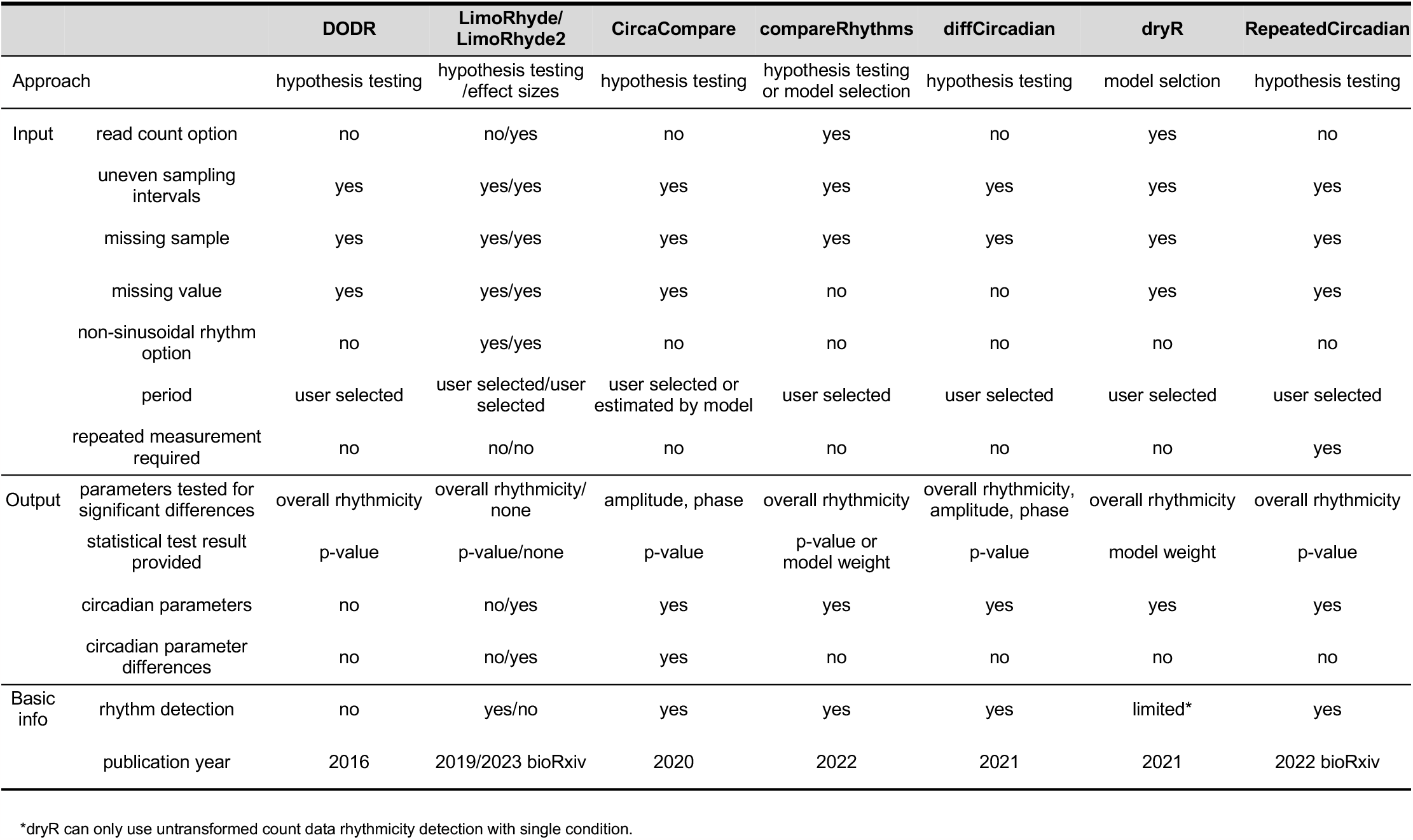
Characteristics of seven algorithms used in this study.

We also compared the output from each algorithm (Table 1). Most algorithms use hypothesis testing appraoch and provide p-values based on the differences in the overall shape of the rhythms (DODR, LimoRhyde, compareRhythms except model selection option, diffCircadian, and RepeatedCircadian), or the difference in specific rhythm parameters, such as amplitude and phase (CircaCompare and diffCircadian). Another approach uses model selection approach (compareRhythms with the model selection option and dryR) and compares how well models with different patterns of rhythmicity (no rhythms, same rhythms, different rhythms) fit the data. Then, it reports the model that best fits the data by using Bayesian Information Criterion (BIC) or Akaike Information Criterion (AIC) which indicates the degree of certainty that the selected model best fits the data. If the best model includes differences in rhythms, the transcript is defined as differentially rhythmic. The compareRhythms package is a collection of separate algorithms, rather than a single approach, and includes options for both model selection and hypothesis testing approaches. Unlike other algorithms, LimoRhyde2 does not perform any statistical tests. Rather, it focuses on quantifying rhythm parameters and effect sizes, including estimating the size of differences between conditions. Thus, this algorithm emphasizes biological relevance over statistical significance. In addition to their statistical test results, most algorithms also return calculated circadian parameters (amplitude and phase) for each condition, while LimoRhyde2 and CircaCompare additionally report calculated differences in parameters between the two conditions.

### Testing algorithm performance with RNA-seq data

To compare these algorithms’ performance side-by-side, we first analyzed the same transcriptomic dataset from mouse fibroblast NIH3T3 cells (9). This dataset consists of two conditions, control and the knockdown of the core circadian clock gene, <u>b</u>rain and <u>m</u>uscle <u>A</u>rnt-like protein 1 (*Bmal1* KD). *Bmal1* is one of the core circadian clock genes, and knockdown or knockout of *Bmal1* dramatically reduces the number of rhythmic transcripts in various mouse tissues and cells (33, 34). For each condition, samples were collected every two hours throughout one circadian cycle (i.e., 24 hours) after serum shock. To analyze these datasets, we selected the model selection option for compareRhythms to directly compare with dryR, which is another algorithm that uses model selection approach. We did not include RepeatedCircadian in the downstream analysis, as it requires data with repeated measurement experimental design.

### Comparison of the performance in detecting rhythmic transcripts

In order to detect differences in rhythmicity under different conditions, transcripts have to be rhythmic in at least one condition *a priori*. Four algorithms (LimoRhyde, CircaCompare, diffCircadian, and dryR [untransformed count data only]) provide the results of a statistical test for the rhythmicity of each transcript under each condition (Table 1: Basic info), while the rhythmicity information provided by compareRhythms depends on which option is selected. The hypothesis testing approaches (“dodr”, “limma”, “voom”, “deseq2”, “edger”, and “cosinor”) report which genes are rhythmic under each condition, although only “dodr” includes a p-value for each condition. For the model selection option, compareRhythms does not provide a separate rhythmicity test, although rhythmicity in each condition can be inferred from which model best fits the data. In contrast, DODR does not include a rhythmicity test, and it is necessary to use other packages, such as MetaCycle (19) and RAIN (20), to pre-identify rhythmic transcripts.

We therefore compared the performance of the four algorithms that provide rhythmicity tests for individual conditions first. Using the control condition samples, we found that CircaCompare detected the highest number of rhythmic transcripts, with 5474, 3630, and 1321 transcripts detected using cutoff values of B.H. q < 0.25, 0.15, and 0.05, respectively (Fig. 1). LimoRhyde detected the second highest number of rhythmic transcripts with the least stringent cutoff (B.H. q < 0.25), followed by dryR and diffCircadian, all of which still detected over 2000 rhythmic transcripts (Fig. 1). With the most stringent cutoff (B.H. q < 0.05), dryR identified more rhythmic transcripts than LimoRhyde and diffCircadian (Fig. 1). As a reference, MetaCycle (19), another algorithm that detects rhythmicity, detected 685, 504, and 297 rhythmic transcripts with B.H. q cutoffs < 0.25, 0.15, and 0.05, respectively. The number of rhythmic transcripts detected by MetaCycle was lower than the number detected by the four algorithms from this study at all three significance thresholds.

**Figure 1.**
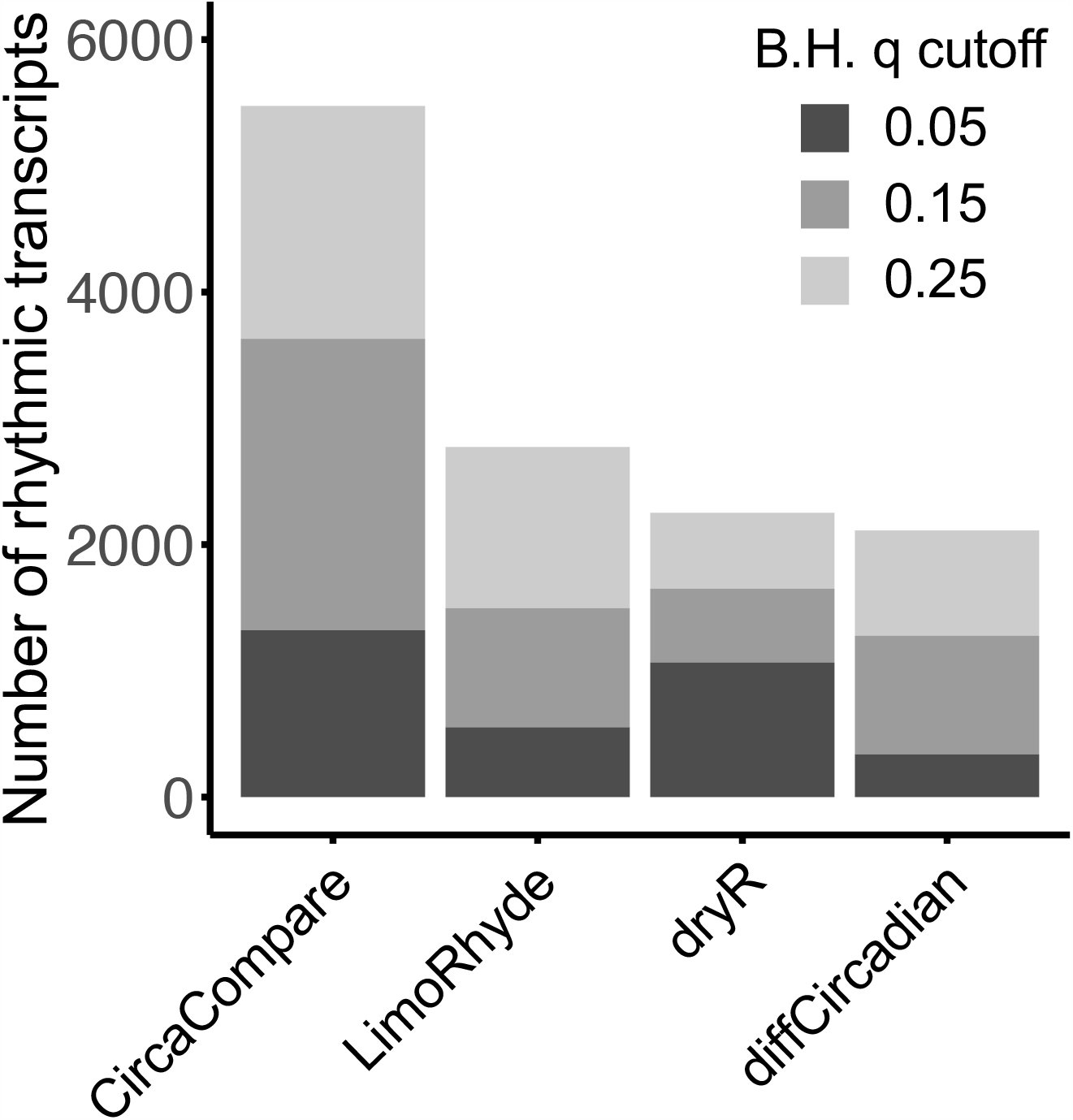
Number of rhythmic transcripts detected by circaCompare, LimoRhyde, dryR, and diffCircadian in the control condition samples. Untransformed reads count were used for dryR, while normalized (RPKM) data were used for circaCompare, LimoRhyde, and diffCircadian.

### Comparison of algorithm performance in detecting differentially rhythmic transcripts

Next, we compared the performance of all the algorithms in detecting differentially rhythmic transcripts (except for LimoRhyde2 which does not perform any statistical tests). Since a transcript must be rhythmic in at least one condition as a pre-requisite for testing for differential rhythmicity, and different algorithms yielded different numbers of rhythmic transcripts (Fig. 1), we used the same set of rhythmic transcripts as an input to directly compare their performance. To this end, we pre-selected 971 transcripts that were detected as rhythmic (B.H. q < 0.25) by MetaCycle in either control or *Bmal1* KD cells (779 in control only, 179 in KD only, and 13 in both). To make comparisons as direct as possible, we also used RPKM values, assumed a 24-hour period, and fit sinusoidal curves for all the algorithms.

Among the algorithms using the hypothesis testing approach and testing for changes in overall rhythmicity, we found that DODR identified more differentially rhythmic transcripts between control and *Bmal1* KD cells than LimoRhyde and diffCircadian with B.H. q cutoffs < 0.25 and 0.15 (Fig. 2A). However, LimoRhyde and diffCircadian detected slightly more transcripts than DODR with the most stringent cutoff at B.H. q < 0.05 (Fig. 2A). For the model selection appraoches that also assess the changes in overall rhythmicity, the recommended BICW threshold to compare two conditions is 0.6 (21, 27). With BICW cutoffs of 0.6, 0.75, and 0.9, dryR identified 434, 286, and 131 transcripts, while compareRhythms identified 407, 272, and 126 transcripts that are differentially rhythmic (Fig. 2B). For CircaCompare, we defined differentially rhythmic transcripts as those that show statistically significant differences (B.H. q < 0.25, 0.15, or 0.05) in either amplitude or phase. Under this condition, CircaCompare identified 304, 229, and 138 differentially rhythmic transcripts, respectively (Fig. 2C). Because of differences in methodology and the necessity of choosing arbitrary cutoff values to define rhythmicity differences, the results cannot be directly compared among these three groups of algorithms.

**Figure 2.**
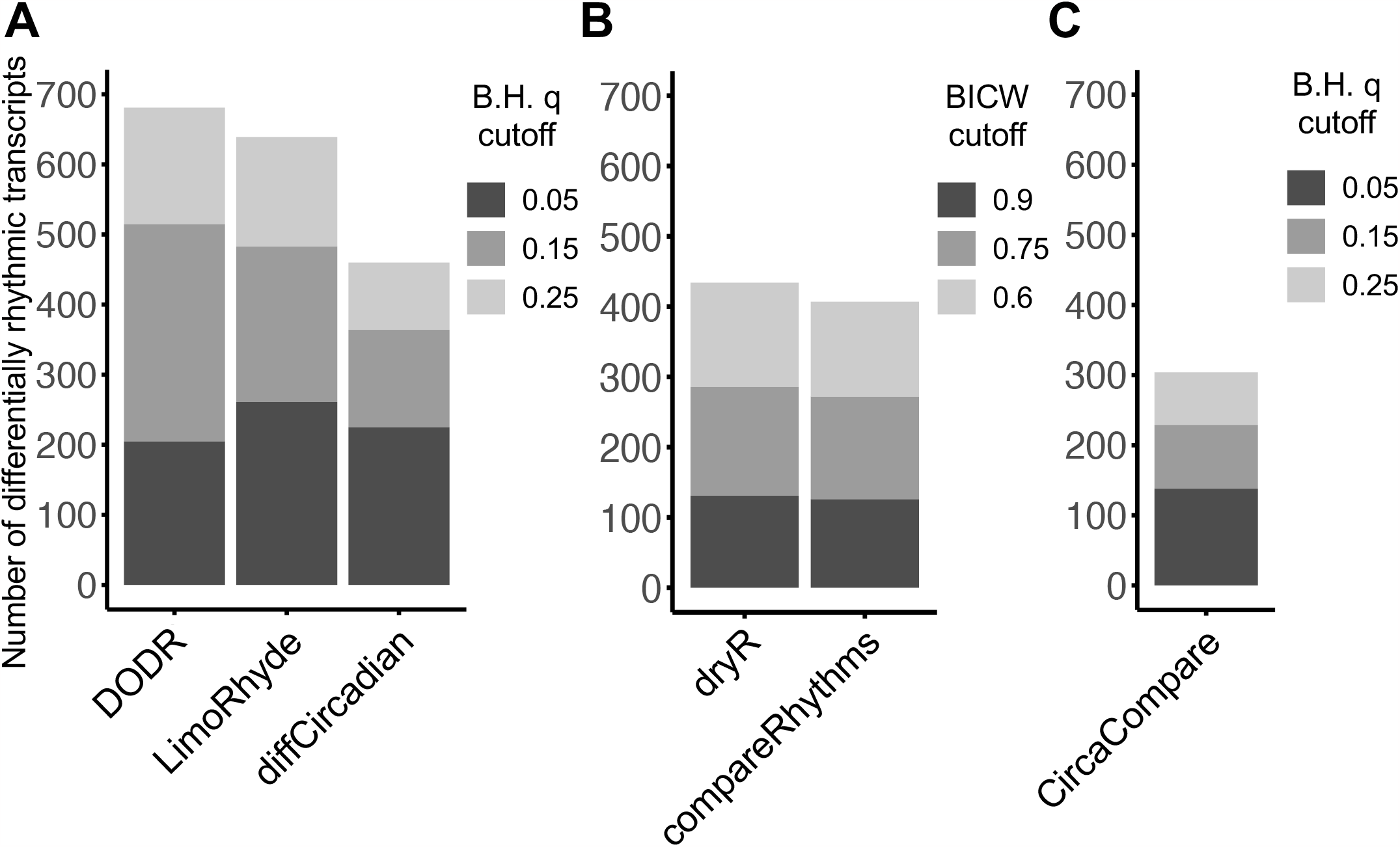
Number of differentially rhythmic transcripts detected by (A) DODR, LimoRhyde, and diffCicadian, (B) dryR and compareRhythms, and (C) circaCompare.

### Overlaps of differentially rhythmic transcripts

Next, we analyzed how many transcripts were commonly detected as differentially rhythmic across different algorithms. Because they perform statistical tests differently, we first focused on the overlaps among the algorithms that are comparable with each other (Fig. 2). Among the algorithms that use hypothesis testing approach and detect overall differential rhythmicity (DODR, LimoRhyde, and diffCircadian), a total of 242 transcripts were commonly detected as differentially rhythmic in all three when using a B.H. q < 0.25 (Fig. 3). More transcripts overlapped between DODR and LimoRhyde, compared to those between DODR and diffCircadian or LimoRhyde and diffCircadian (Fig. 3B). We also compared the algorithms based on the model selection approach (dryR and compareRhythms) with BICW > 0.6 as a cutoff. We found that 407 differentially rhythmic transcripts identified by compareRhythms were also detected by dryR, while dryR detected an additional 27 transcripts as differentially rhythmic. We also compared the overlaps among the six algorithms, even though these results are not technically comparable. We found that 85 transcripts were commonly identified as differentially rhythmic by all six algorithms (Fig. 4).

**Figure 3.**
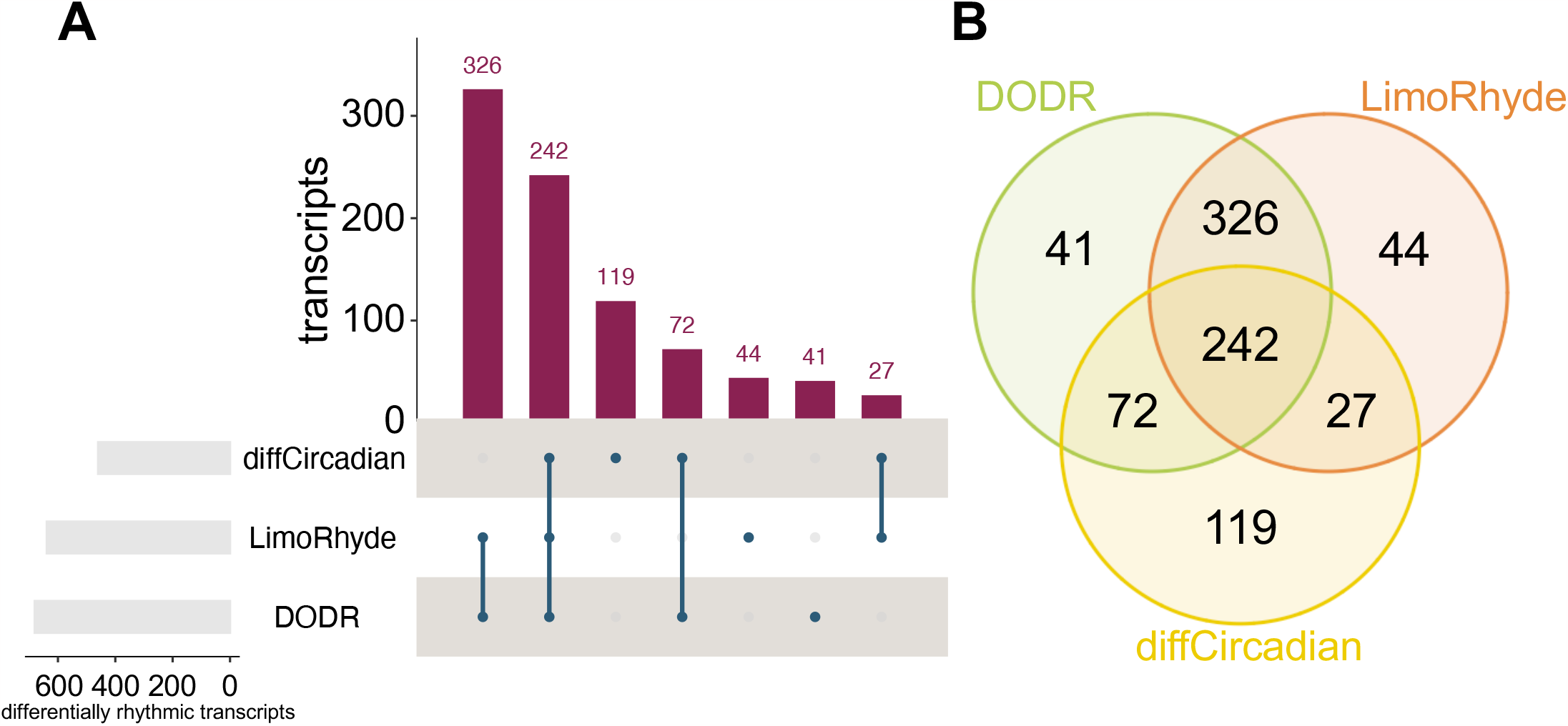
The number of overlapping transcripts identified as differentially rhythmic among diffCircadian, LimoRhyde, and DODR with B.H. q cutoff of 0.25. (A) Upset plot. Each intersection is exclusive (i.e. each transcript can appear in only one intersection). (B) Venn diagram.

**Figure 4.**
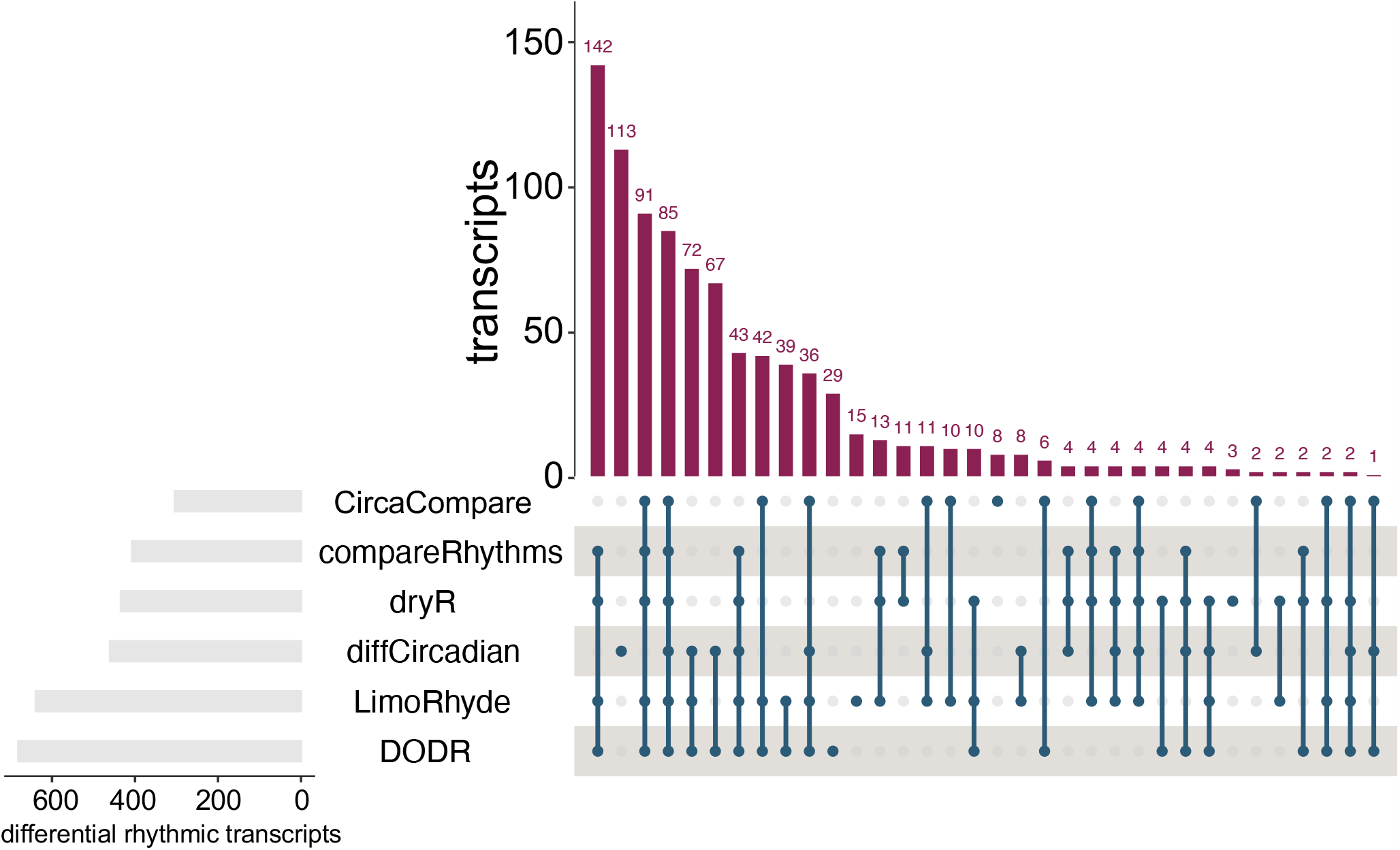
The number of overlapping transcripts identified as differentially rhythmic among six the algorithms (B.H. q < 0.25 or BICW >0.6). Each intersection is exclusive (i.e. each transcript can appear in only one intersection).

### Are differences in amplitude and phase detected by each algorithm similar?

Five algorithms (LimoRhyde2, CircaCompare, compareRhythms, diffCircadian, and dryR) also report circadian parameters (i.e., phase and amplitude) for each transcript (Table 1), allowing us to compare the differences in these parameters between conditions. Because compareRhythms, diffCircadian, and dryR only calculate each parameter for each condition, we manually calculated the difference in phase or amplitude by subtracting the value for the control condition from that of the *Bmal1* KD condition. We then compared these values with the differences calculated directly by CircaCompare and LimoRhyde2. We found 32 transcripts that were detected as rhythmic under both conditions and commonly identified as differentially rhythmic by all five algorithms with the least stringent cutoff (i.e., B.H. q < 0.25 or BICW > 0.6; LimoRhyde significance test results were used in place of LimoRhyde2) (Fig. 5A, B). To assess the similarities in the changes of phase and amplitude, we conducted a pairwise correlation analysis of their respective outputs (Fig. 5C, D). We found that the phase and amplitude differences measured by all five algorithms are positively correlated with each other (Pearson: p < 4.3 x 10^-7^) (Fig. 5C, D). The correlation is particularly strong (correlation coefficient = 1) among CircaCompare, compareRhythms, diffCircadian, and dryR for both phase and amplitude. In fact, the differences in phase and amplitude are identical between CircaCompare and diffCircadian, as well as between compareRhythms and dryR. These indicate that the changes in phase and amplitude measured by these five algorithms were remarkably similar.

**Figure 5.**
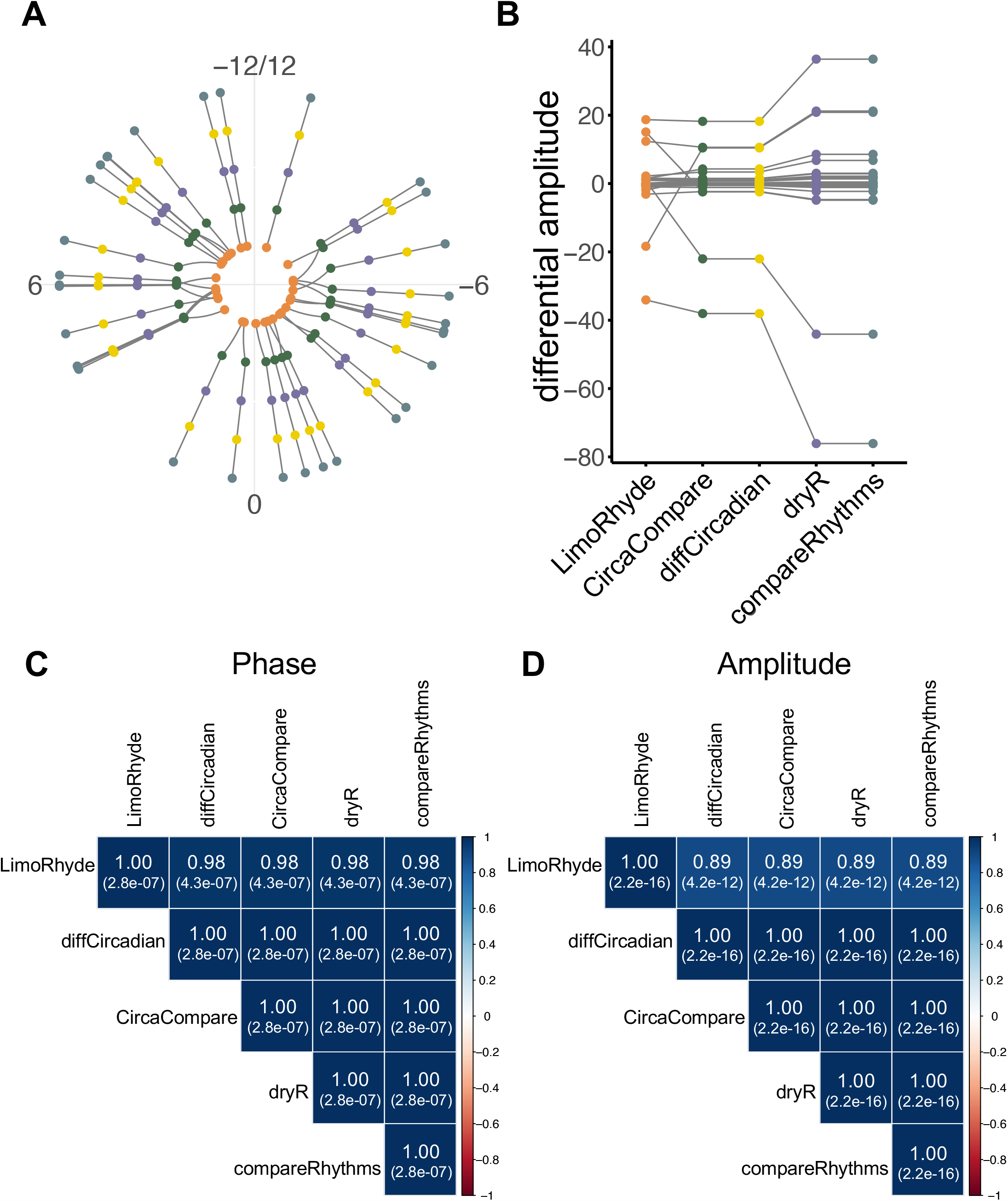
Comparison of the changes in phase or amplitude of differentially rhythmic transcripts. Differences in (A) phase or (B) amplitude detected by five algorithms. Orange: LimoRhyde2, Green: circaCompare, Purple: dryR, Yellow: diffCircadian, and Blue: compareRhythms. Each dot represents a measurement for a transcript from one of the algorithms, and each line represents each transcript. (C, D) Heatmap displaying pairwise Pearson correlation analysis of the differences in (C) phase or (D) amplitude calculated by the five algorithms. Blue and red denoted positive and negative correlation, respectively. The correlation coefficient and p-value (number in brackets) are shown in each box.

## Discussion

In this study, we compared seven algorithms designed to detect differential rhythmicity between transcriptomic datasets. These algorithms show distinct features and use different approaches in defining the rhythmicity of transcripts as well as in identifying differences in rhythmicity between different conditions (Table 1). They also report different numbers of rhythmic transcripts and differentially rhythmic transcripts even with the same statistical threshold (Figs. 1, 2). Nevertheless, we still detected large overlaps in the sets of differentially rhythmic transcripts detected by each algorithm (Figs. 3, 4), and the phase and amplitude differences measured by each algorithm for the set of commonly detected transcripts also showed high similarities (Fig. 5).

These algorithms provide important and substantial advancements over Venn Diagram Analysis. However, they also face several key limitations. Successfully detecting a change in rhythmicity depends heavily on statistical power, and this is affected by the amount of noise in the data, the magnitude of the change, and the number of data points (e.g., number of sampling time points, number of replicates) (27). Different studies with different amounts of data noise or different experiment designs can have large differences in statistical power, potentially resulting in large differences in the number of differentially rhythmic transcripts that are detected (35, 36). Another important consideration is that statistical significance does not always indicate biological significance. LimoRhyde2 attempts to address this by focusing on estimating rhythm parameters and effect sizes, and the magnitude of rhythm differences. However, establishing a direct link between the amplitude (or change in amplitude) of a rhythm and its biological significance remains challenging. Follow-up work will likely be necessary to uncover if differences in rhythms are biologically relevant. Finally, these algorithms are only designed to compare rhythms between datasets that share the same scale. Data with different units or scales, for instance, comparing rhythms between transcriptomic data and proteomics data, or between transcriptomic data and RNA synthesis or degradation rates (9), cannot be effectively analyzed using these algorithms. VDA analysis, in this case, can be used as it evaluates rhythmicity under each condition independently. However, it still has the issues of indirect analysis, uncontrolled FDRs, and a lack of sensitivity to quantitative circadian changes. The development of algorithms capable of detecting differential rhythmicity in such datasets would represent a valuable and much-needed contribution to the field.

As users, we found several features helpful when analyzing data and recommend that developers include them in their future algorithms and R packages. First, we prefer algorithms that exhibit wide applicability and have the flexibility to accept a wide range of data types and experiment designs, as this would benefit a diverse range of users. Second, we advocate for a complete and detailed description of the algorithm, presented in a way that non-statisticians can understand what the algorithm does and interpret the results correctly. For instance, clearly describe the hypotheses being tested by a statistical test, or the steps used for model fitting. Thorough code documentation and “vignettes” with worked examples are also important for users learning how to run the analysis. Third, it is important that algorithms offer explicit and comprehensive information regarding the input data requirements, including formatting requirements and any other characteristics (e.g., Do the data need to have Gaussian-distributed residuals?). As for output, it is preferable for algorithms to provide calculated statistical values (e.g., p-values or BICW) rather than applying statistical threshold filtering within the code and only returning qualitative results. Finally, algorithms that allow users to specify the output they require have a broader applicability. This flexibility is particularly useful when dealing with large datasets or computationally intensive tasks, as it can significantly reduce both time and computational load or memory requirements.

In addition to transcriptomic data, a wide variety of circadian time series data should be analyzable by one or more of these approaches as long as the data sufficiently meet the assumptions of the underlying regression model and other algorithm components. In addition to the requirements listed in Table 1, non-transcriptomic data can be analyzed with any of the algorithms in this study if they have residuals (the difference between each data point’s actual value and the regression model’s predicted value) with a Gaussian distribution and constant variance and lack problematic outlier data points. The only exception is RepeatedCircadian, which should only be used for repeated measurement experiments. For input data featuring non-Gaussian distributions or outliers, there are several options. For example, DODR can accommodate data with non-Gaussian distributions as long as the distribution is the same between the two groups being compared (22). Positive integer count data that have similar characteristics to RNA-seq read counts (negative binomial residual distribution and variance that changes with the mean of the data) can be analyzed with LimoRhyde2, compareRhythms, and dryR. However, in this case, users should select an option designed for RNA-seq read count data (e.g., voom, DESeq2, or edgeR based method) (21, 25, 27). Alternatively, data can potentially be transformed before analysis so that the residuals have a Gaussian distribution with constant variance, and then analyzed using one of the Gaussian-assuming approaches listed above (23). If there are outliers in the data, DODR can be used (22). RNA-seq datasets contain data on thousands of features (e.g., thousands of transcripts), yet it is common for other types of circadian studies to measure only one or a few features. In this case, DODR, CircaCompare, dryR (drylm), and RepeatedCircadian are the most straightforward to use.

Overall, we hope our study will help future users in selecting the algorithm that is most suitable for their data and specific research goals. Understanding the characteristics of these algorithms is a critical first step towards this, and thus our analysis will contribute to improved analyses of differential rhythmicity in transcriptomic data.

## Acknowledgment

We thank the members of the Kojima lab for manuscript proofreading. This work was supported by National Institute of Health F31 AG071393 (to B.A.U) and R01 GM126223 (to S.K.).

## Data Availability

There are no primary data in the paper.

